# Characterisation of Antibody Interactions with the G Protein of Vesicular Stomatitis Virus Indiana Strain and Other Vesiculovirus G Proteins

**DOI:** 10.1101/330910

**Authors:** Altar M Munis, Maha Tijani, Mark Hassall, Giada Mattiuzzo, Mary K Collins, Yasuhiro Takeuchi

## Abstract

Vesicular stomatitis virus Indiana strain G protein (VSVind.G) is the most commonly used envelope glycoprotein to pseudotype lentiviral vectors (LV) for experimental and clinical applications. Recently, G proteins derived from other vesiculoviruses (VesG), for example Cocal virus, have been proposed as alternative LV envelopes with possible advantages compared to VSVind.G. Well-characterised antibodies that recognise VesG will be useful for vesiculovirus research, development of G protein-containing advanced therapy medicinal products (ATMPs), and deployment of VSVind-based vaccine vectors. Here we show that one commercially available monoclonal antibody, 8G5F11, binds to and neutralises G proteins from three strains of VSV as well as Cocal, and Maraba viruses, whereas the other commercially available monoclonal anti-VSVind.G antibody, IE9F9, binds to and neutralises only VSVind.G. Using a combination of G protein chimeras and site-directed mutations, we mapped the binding epitopes of IE9F9 and 8G5F11 on VSVind.G. IE9F9 binds close to the receptor binding site and competes with soluble low-density lipoprotein receptor (LDLR) for binding to VSVind.G, explaining its mechanism of neutralisation. In contrast, 8G5F11 binds close to a region known to undergo conformational changes when the G protein moves to its post-fusion structure, and we propose that 8G5F11 cross-neutralises VesGs by inhibiting this.

**IMPORTANCE:** VSVind.G is currently regarded as the gold-standard envelope to pseudotype lentiviral vectors. However, recently other G proteins derived from vesiculoviruses have been proposed as alternative envelopes. Here, we investigated two anti-VSVind.G monoclonal antibodies for their ability to cross-react with other vesiculovirus G proteins, and identified the epitopes they recognise, and explored the mechanisms behind their neutralisation activity. Understanding how cross-neutralising antibodies interact with other G proteins may be of interest in the context of host-pathogen interaction and co-evolution as well as providing the opportunity to modify the G proteins and improve G protein-containing medicinal products and vaccine vectors.

## INTRODUCTION

The rhabdovirus, vesicular stomatitis virus Indiana stain (VSVind), has been used ubiquitously as a model system to study humoral and cellular immune responses in addition to being a promising virus for oncolytic virotherapy against cancer (1-3). Furthermore, its single envelope G protein (VSVind.G) is the most commonly used envelope to pseudotype lentiviral vectors and serves as the gold-standard in many experimental and clinical studies (4-6). Both receptor recognition and membrane fusion of the wild-type virus, as well as the pseudotyped particles, are mediated by this single transmembrane viral glycoprotein that homotrimerises and protrudes from the viral surface (7-9). Recently G proteins derived from other vesiculovirus subfamily members, namely, Cocal, Piry, and Chandipura viruses, have been proposed as alternative envelopes for lentiviral vector production due to some possible advantages over VSVind.G (10-12).

Although some antigenic and biochemical characteristics of VSVind.G have been reported (1, 7, 13-20), there is still little known about the other vesiculovirus G proteins (VesG) and there is a general lack of reagents commercially available to identify, detect, and characterise them. In the past, monoclonal antibodies (mAbs) have been used to extensively study the antigenic determinants found on viral glycoproteins, e.g. hemagglutinin (HA) of influenza virus, the gp70 protein of murine leukaemia virus (MLV), and rabies virus G protein (21-25). These previous studies, especially on the influenza virus strains and the rabies virus have led to invaluable findings on the structure and function of the glycoproteins allowing identification of epitopes essential in virus neutralisation (25-27). In addition, mAbs have proven useful in viral pathogenesis studies as mutants selected by antibodies, in many cases demonstrated altered pathogenicity to their wild-type counterparts (28-30). Therefore, identification of antibodies that recognise VesG will not only be extremely valuable for vesiculovirus research but also aid in the development of G protein-containing advanced therapy medicinal products (ATMP) and vaccine vectors.

Here we show two anti-VSVind.G antibodies, 8G5F11 and a goat polyclonal antibody, VSV-Poly (31, 32), can cross-react with a variety of the VesG and cross-neutralise VesG-LV. We also demonstrate that the other commercially available extracellular monoclonal anti-VSVind.G antibody IE9F9 lacks this cross-reactivity. We further characterise the two mAbs, 8G5F11 and IE9F9, with regards to their relative affinities towards various VesG, binding epitopes, and cross-neutralisation strengths.

## RESULTS

### Investigation of antibody cross-reactivity with VesG

To investigate antibody binding to different vesiculovirus envelope glycoproteins (G proteins), we prepared plasmid pMD2-based vectors expressing six different vesiculovirus G proteins (VesG): VSVind.G, Cocal virus G (COCV.G), Vesicular stomatitis virus New Jersey strain G (VSVnj.G), Piry virus G (PIRYV.G), Vesicular stomatitis virus Alagoas strain G (VSVala.G), and Maraba virus G (MARAV.G) (Figure 1A). HEK293T cells were transfected with these plasmid constructs, stained with the different antibodies, and analysed via flow cytometry. While IE9F9 only bound to VSVind.G, anti-VSVind.G monoclonal antibody 8G5F11 and VSV-Poly both could recognise various VesG with varying binding strengths (Figure 1B). PIRYV.G, the most distant vesiculovirus G investigated with approximately 40% identity to VSVind.G on amino acid level, could be recognised by VSV-Poly while 8G5F11 did not bind to it.

**Figure 1:**
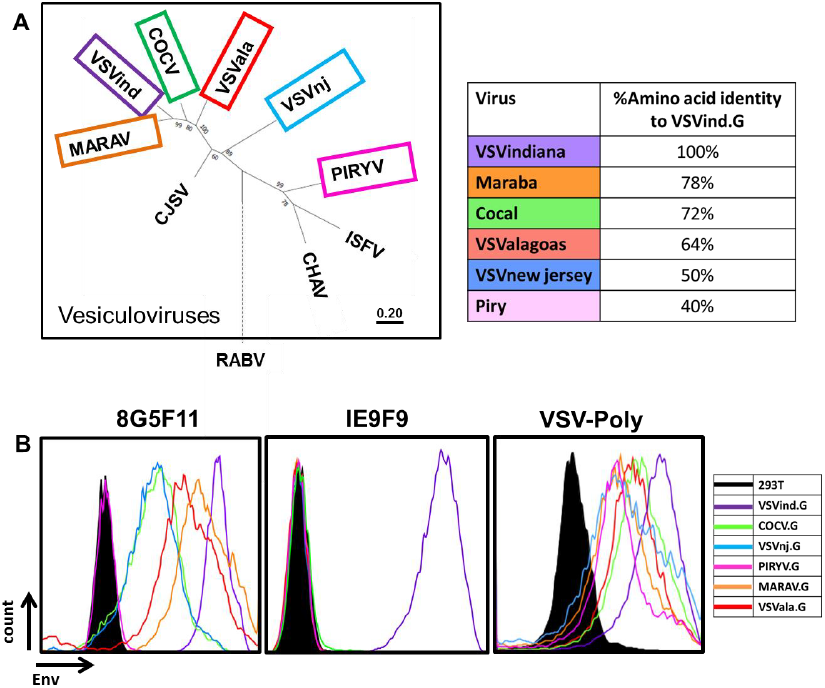
8G5F11 and VSV-Poly cross-react with a variety of VesG while IE9F9 only binds to VSVind.G. **(A)** G proteins of the major vesiculoviruses, as well as the G protein of the rabies virus (RABV), were analysed with regards to their phylogenetic relationship. The tree amongst VesG is drawn to scale, with branch lengths measured in the number of substitutions per site, depicted in the linear scale. VSVind: Vesicular stomatitis virus Indiana strain, COCV: Cocal virus, VSVnj: Vesicular stomatitis virus New Jersey strain, PIRYV: Piry virus, CJSV: Carajas virus, CHAV: Chandipura virus, ISFV: Isfahan virus, MARAV: Maraba virus, VSVala: Vesicular stomatitis virus Alagoas strain. Vesiculoviruses that we investigated are highlighted in boxes and percentage amino acid identities to VSVind.G are summarised in the table on the right-hand side.**(B)** Histograms represent the binding of the antibodies to the VesG expressed on the surface of transfected HEK293T cells. The strength of cross-reaction is depicted via the different MFIs of the histograms. On the other hand, IE9F9 only bound to VSVind.G. Data shown is one of the three repeats performed.

### Characterisation of IE9F9 binding, 8G5F11 cross-reactivity and its affinity towards other VesG

To confirm that the difference of 8G5F11 binding to VesG was indicative of the mAb affinity towards VesG and not a difference in relative expression levels of the G proteins, we synthesised chimeric G proteins. The endogenous transmembrane and C-terminal domains of VesG were switched with that of VSVind.G (Figure 2A). Following the expression of these chimeric G proteins in HEK293T cells, we investigated 8G5F11 and IE9F9 binding saturation using quantitative flow cytometry while the relative expression levels of the G proteins were monitored using an intracellular anti-VSVind.G mAb, P5D4 (Figure 2B). 8G5F11 showed a wide range of affinities towards VesG: while its affinity for MARAV.G was comparable to that of VSVind.G, its interactions with COCV.G and VSVnj.G were much weaker.

**Figure 2:**
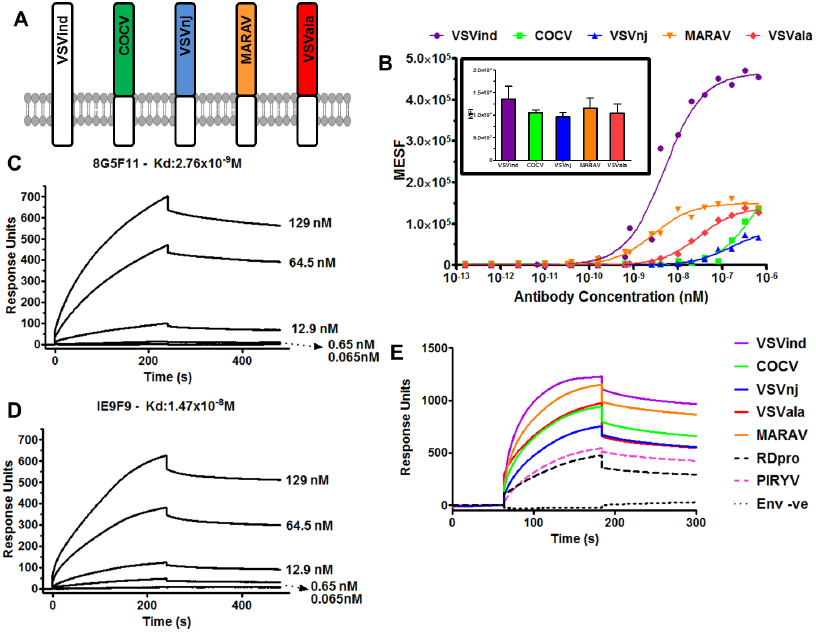
Investigation of 8G5F11 and IE9F9 affinities towards VSVind.G and characterisation of 8G5F11 cross-reactivity. **(A)** Schematic representation of the chimeric vesiculovirus G proteins with VSVind.G transmembrane and C-terminal domains. **(B)** HEK293T cells expressing chimeric VesG were incubated with serial dilutions of 8G5F11 and analysed via flow cytometry. MFIs of the fluorescent signals were converted into the number of fluorophores using the MESF standard curve according to manufacturer’s instructions, the background signal from mock-transfected HEK293Ts was subtracted and binding saturation curves were plotted. The varying affinity of the mAb towards different VesG is demonstrated by the shift in the slope of the binding curves. The curves were fitted, and dissociation constants (Kd) calculated using the software GraphPad Prism 5 modelling the interaction as 1:1 specific binding: VSVind.G: 2.64×10^−9^M, COCV.G: 5.88×10^−7^M, VSVnj.G: 1.57×10^−7^M, MARAV.G: 4.13×10^−9^M, VSVala.G: 3.09×10^−9^M. Data shown represent the mean of three repeats performed in duplicates. **(inset)** The expression levels of the chimeric G proteins were determined via intracellular P5D4 staining. Data shown represent the mean +/−SD of three repeats performed in duplicates. Surface plasmon resonance (SPR) analysis of **(C)** 8G5F11 and **(D)** IE9F9 binding to immobilized Gth in HBS-EP buffer. **(E)** Surface plasmon resonance analysis of Ves.G-LV (1×10^8^ TU/ml) binding to immobilized 8G5F11 in HBS-EP buffer. The binding curves are normalised with regards to the relative response of unenveloped LV particles (Env-ve) which is regarded as the background. SPR data shown is one of the three repeats performed.

To consolidate this finding, we further investigated these mAb-G protein interactions via surface plasmon resonance. First, to quantify mAb binding to G protein monomers under conformationally correct folding, we immobilised wild-type (wt) VSVind.G produced by thermolysin limited proteolysis of viral particles (Gth) (7, 17) and tested the dose-dependent binding of the two mAbs (Figure 2C-D). The measured Kd values for 8G5F11 and IE9F9 binding to VSVind.G were 2.76nM and 14.7nM respectively. To further analyse the VesG-8G5F11 interaction we immobilised the mAb and investigated VesG pseudotyped lentiviral vector (LV) binding. Since pseudotyped LV particles contain many trimeric G protein spikes (33), the analysis of the interaction between VesG binding to immobilised 8G5F11 reflects avidity. Dose-response binding of VSVind.G resulted in a strong response implying high avidity. (Supplementary Figure S1). When identical doses of VesG-LV at 1×10^8^ TU/ml were injected on immobilized 8G5F11, similar patterns of binding were observed to that of quantitative flow cytometry, in the order of strength of VSVind > MARAV > VSVala > Cocal > VSVnj (Figure 2E). Unrelated RDpro envelope pseudotyped LVs were utilised as negative control to deduce unspecific interaction of enveloped particles with immobilised mAb. PIRYV.G-LV demonstrated a similar response to that of RDpro-LV indicative of the lack of binding between the G protein and 8G5F11.

### Determining the cross-neutralisation abilities of anti-VSVind.G antibodies

These three antibodies were evaluated for their ability to neutralise VSVind.G and VesG pseudotyped LVs (Figure 3). 8G5F11 demonstrated varying strengths of neutralisation against VesG pseudotyped LVs, IC50 values ranging from 11.5ng/ml to 86.9μg/ml (Figure 3A). There was however limited correlation between G proteins’ binding strength and sensitivity of LV, e.g. VSVnj.G-LV was more sensitive than COCV.G-LV (Figure 3A) while COCV.G binding was stronger (Figure 1 and 2). IE9F9 neutralised only VSVind.G-LV at 137ng/ml IC50, about 12-fold weaker than 8G5F11 (Figure 3B). In the case of VSV-Poly, we only observed cross neutralisation at high serum concentrations (Figure 3C). Furthermore, although VSV-Poly bound to PIRYV.G, it did not neutralise PIRYV.G-LVs.

**Figure 3:**
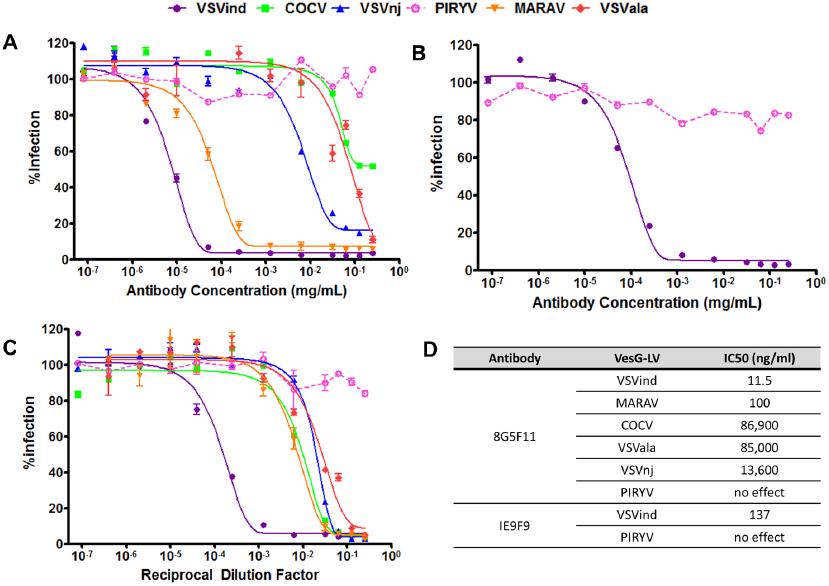
Neutralisation activity of mAbs and VSV-Poly. Neutralisation of VesG-LV by **(A)** 8G5F11, **(B)** IE9F9, and **(C)** VSV-Poly. Solid lines signify the neutralisation effect observed while the dotted lines indicate the lack of neutralisation. **(D)** Calculated IC50 values for 8G5F11 and IE9F9, depicting the potency of neutralisation. The curves were fitted using the software GraphPad Prism 5 modelled as an [inhibitor] vs. response curve with variable Hill Slopes and IC50 values calculated. Data shown represent the mean +/−SD of three repeats.

### Mapping the epitopes of anti-VSVind.G mAbs and identification of key amino acid residues that dictate antibody binding and neutralisation

To map where the neutralising antibodies might bind to on the G protein surface a series of chimeric G proteins between VSVind.G and COCV.G were constructed. The initial binding and neutralisation studies performed with these chimeras enabled us to narrow down the epitopes of these mAbs to lie between amino acid (aa) residues 137-369^1^ on VSVind.G (Supplementary Figure S2). Furthermore, looking at previously published data on 8G5F11 and IE9F9’s epitopes obtained through mutant virus escape assays (1, 13-15) we concentrated on two distinct regions on VSVind.G and synthesized 22 different mutant G proteins to study the epitopes (Figure 4). The mutants were cloned into the pMD2 backbone and their functionality were investigated via LV infection and antibody binding assays. All G proteins were confirmed to be functional and could successfully pseudotype LVs yielding comparable titres to their wild-type (wt) counterparts. Furthermore, their relative expression levels were monitored by intracellular P5D4 which also recognises the intracellular domain of COCV.G. Lastly, they could be detected by extracellular VSV-Poly implying there weren’t any substantial protein display issues (Supplementary Figure S3).

**Figure 4:**
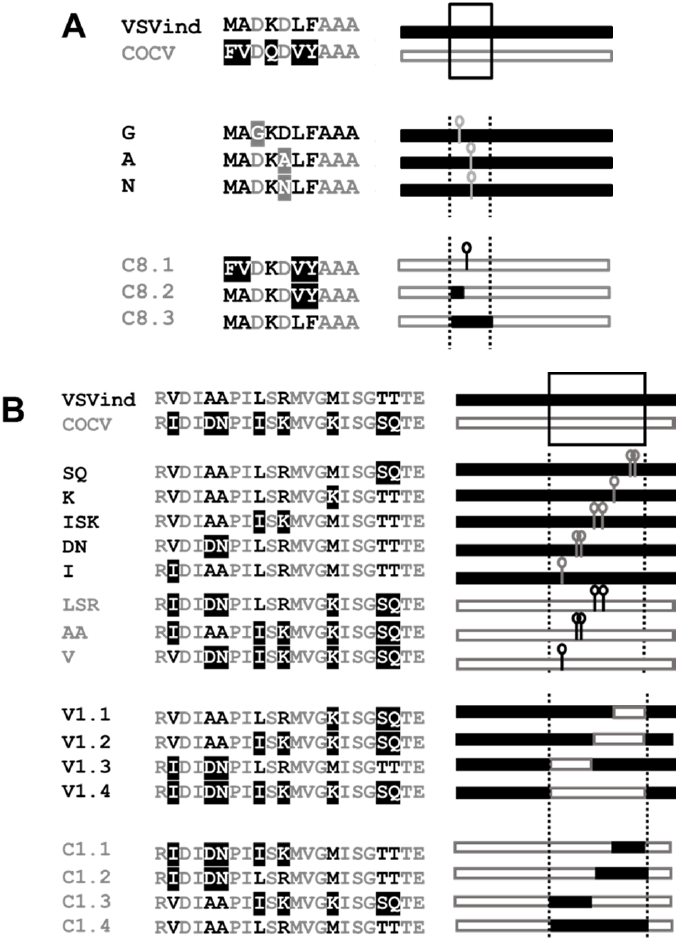
Mutants and chimeric G proteins produced for epitope mapping. Mutants and chimeras produced for epitope mapping of monoclonal antibodies **(A)** 8G5F11 and **(B)** IE9F9. Names and linear representations of the mutants and chimeras are listed on either side of the amino acid alignments of the regions where mutations were made. Amino acid alignment legend: Black; residues from wt VSVind.G, white with black background; residues from wt COCV.G, grey; shared residues, white with grey background; previously identified mutants (15). Linear G protein representations: the regions that the mutations were carried out at are represented by dotted lines. Black bars represent wt VSVind.G sequences while grey-bordered bars are for wt COCV.G residues. Point mutations are denoted by a bar and a circle.

We first investigated antibody binding to these G proteins via flow cytometry. Relative expression levels of the mutants were determined by extracellular VSV-Poly and intracellular P5D4 stains. For both sets the relative difference between expression levels of mutant and wt proteins was in most cases less than two-fold (Figure 5A-B). In the case of 8G5F11, binding to VSVind.G mutants was reduced by approximately 100-fold while the changes on COCV.G enabled these mutants to bind to 8G5F11 at similar levels to that of wt VSVind.G (Figure 5C). This change in binding could also be observed on a western blot. While none of the VSVind.G mutants could be visualized, 8G5F11 could bind to COCV.G chimera C8.3 (data not shown). It can be inferred from these results that aa 257-259 (DKD) are the main residues that dictate 8G5F11 binding to G proteins.

**Figure 5:**
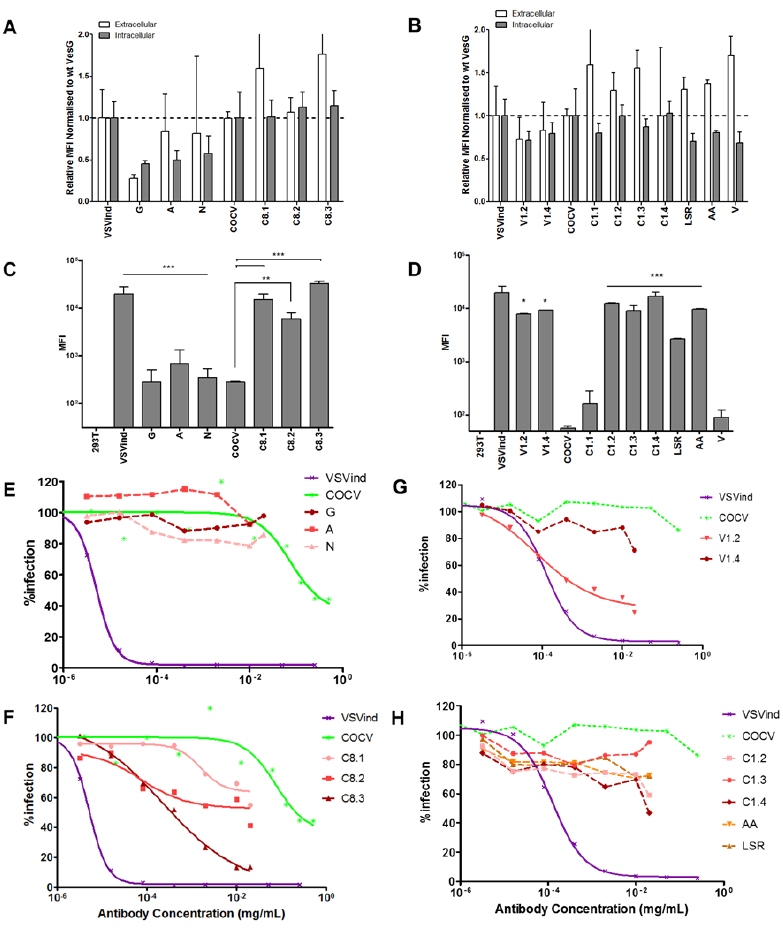
Investigation of antibody binding to mutant G proteins and neutralisation of mutant-LVs. HEK293T cells were transfected to express the mutant G proteins on their surface. **(A-B)** The cells expressing chimeric mutants were stained with extracellular VSV-Poly (white bars) and intracellular P5D4 (grey bars) as expression control for the G proteins. The measured MFI values were normalised to the wt VesG signals for each set of mutants. The same population of cells were also incubated with **(C)** 8G5F11 and **(D)** IE9F9 at saturating concentrations. One-way ANOVA analysis with Dunnett’s post-test was performed to compare the MFI values of mutant G proteins to that of their wild-type counterpart. Legged lines denote the significance of a single comparison, while straight lines signify all the individual comparisons within the group share the denoted significance unless otherwise stated (*, p<0.05; **, p<0.01; ***, p<0.001). This assay was performed three times in duplicates; mean +/−SD is plotted above. The neutralisation curves for select mutant and chimeric G pseudotyped LVs are plotted for **(E-F)** 8G5F11 and **(G-H)** IE9F9. Solid lines signify the neutralisation effect observed. **(E-G)** Previously reported reductions in binding for VSVind.G mutants translated into either complete or partial resistance to neutralisation by both antibodies. For COCV.G mutants **(F-H),** the mutations conferred the G proteins sensitivity to neutralisation by 8G5F11 but not by IE9F9. The curves were fitted using the software GraphPad Prism 5 modelled as an [inhibitor] vs. response curve with variable Hill Slopes. Data shown represent the mean from three experiments performed in independent triplicates.

On the other hand, for IE9F9 no statistically significant changes in antibody binding were observed for VSVind.G mutants (data not shown) except for chimeras V1.2 and V1.4 (Figure 5D). However, there was a substantial gain of binding effect for COCV.G mutants. While IE9F9 doesn’t bind to wt COCV.G, mutations of amino acid residues LSR and AA (Figure 4) alone led to significant increase in the fluorescence signal, thus antibody binding, C1.4 with both LSR and AA had a comparable MFI level to that of wt VSVind.G.

Neutralisation profile of both VSVind.G and COCV.G mutants was also examined (Figure 5E-H). While LVs pseudotyped with VSVind.G mutants were not neutralised (Figure 5E), varying degrees of sensitivity were observed for COCV.G mutants with the strongest binder being the most sensitive (Figure 5F). On the other hand, this was not the case for IE9F9 mutants. While dose-dependent neutralisation of V1.2-LV was observed, VSVind.G mutant V1.4-LV was resistant to IE9F9 neutralisation (Figure 5G). Furthermore, no effect was observed on COCV.G mutant LV infection even though all bound to the mAb, some at similar levels to wt VSVind.G (Figure 5H). The data shows that while 8G5F11 employs a neutralisation mechanism that is universally effective amongst the tested VesG, IE9F9’s is VSVind.G specific and binding does not necessarily result in neutralisation.

### Investigation of neutralisation mechanisms utilised by the mAbs: binding competition with low-density lipoprotein receptor (LDLR)

Antibodies neutralise viruses and viral vectors by several mechanisms. Many neutralising antibodies (NAbs) prevent virions from interacting with cellular receptors (34). VSVind.G’s major receptor has been identified as the low-density lipoprotein receptor (LDLR) (33, 35). Therefore, we investigated the binding competition between 8G5F11 and IE9F9 with LDLR via SPR as a potential neutralisation mechanism for the mAbs (Figure 6). Gth immobilised on the chip surface was saturated with repeated injections of 8G5F11 and IE9F9. This was followed by an injection of recombinant soluble human LDLR (sLDLR) and its binding to Gth was examined. While sLDLR was able to bind to Gth following 8G5F11 saturation as well as Gth without antibody exposure (buffer control), this interaction was almost completely abrogated by IE9F9. These data suggest that IE9F9, but not 8G5F11, neutralises VSVind.G-LV by blocking the G protein-receptor interaction either through steric hindrance or direct competition.

**Figure 6:**
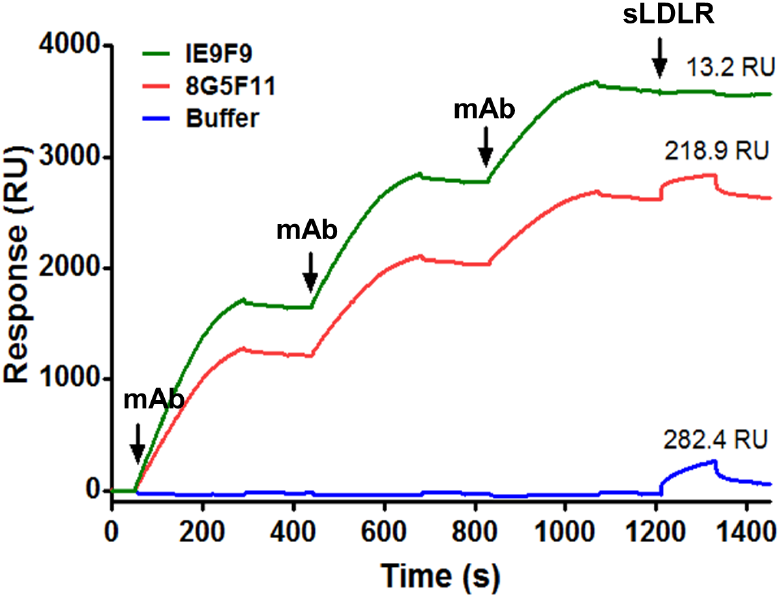
IE9F9 hinders sLDLR binding to Gth. 8G5F11 and IE9F9 were injected over immobilised Gth at 10μg/ml concentration three times to achieve binding saturation. Following this, sLDLR was injected over the chip at a concentration of 10μg/ml and its binding to Gth was measured. As buffer control an identical sLDLR injection was performed following multiple injections of HBS-EP running buffer. Measured sLDLR binding levels are indicated above the binding response curves and times of injections are marked with arrows. The data presented represent one of the three repeats performed.

## DISCUSSION

VSVind.G is the most commonly used envelope glycoprotein to pseudotype LVs for experimental and clinical applications. VSVind.G pseudotyped LVs can be produced in high titres and can infect a range of target cells. However, VSVind.G is cytotoxic to cells; thus, it is difficult to express it constitutively (36, 37). Moreover, VSVind.G pseudotyped LVs can be inactivated by human serum complement which limits their potential *in vivo* use (38-42). Therefore, there is a clear need for alternative envelopes to pseudotype LVs. Some of the most recent alternative envelopes that have been utilised are the G proteins of the other vesiculovirus family members (10-12). However, one drawback of using these new G proteins is that there are no reagents commercially available to identify or characterise them.

In this report, we have demonstrated that 8G5F11 monoclonal antibody can, unlike VSVind.G specific IE9F9, cross-react with a variety of the VesG and cross-neutralise VesG-LV. Furthermore, we characterised a goat anti-VSVind.G polyclonal antibody which also can bind and neutralise a wide range of vesiculovirus G proteins.

The cross-reactive monoclonal 8G5F11 demonstrated interesting characteristics. Its high cross-reactivity even towards more distant relatives of VSVind.G such as VSVnj.G suggested that it might be recognising a well-conserved epitope. However, the results of the binding saturation assay didn’t correlate with phylogenetic relativity. It revealed that its affinity towards COCV.G, one of the closest relatives of VSVind.G, was one of the weakest amongst the VesG investigated with almost a 250-fold difference compared to VSVind.G (Figure 2B).

This discrepancy can be explained through fine mapping of the 8G5F11 epitope. We identified the amino acids 257-259, DKD, as the key residues on VSVind.G for 8G5F11 binding. On VSVind.G the two negatively charged aspartic acid residues flank the positively charged lysine possibly contributing towards the structure of the α-helix form through salt-bridges (7, 16, 17). When either of the aspartic acid residues is mutated to a neutral residue a significant reduction in binding is observed. When this is compared to the corresponding three residues on other VesG, the antibody binding is dependent on the overall charge of these three residues rather than the ones surrounding them. In MARAV.G, these residues are identical to VSVind.G, explaining why the antibody has similar strength of binding to these two G proteins (Supplementary Figure S4). On the other hand, VSVala.G binds 8G5F11 with high affinity although these residues are not fully conserved, as in VSVala.G the second aspartic acid residue is replaced with a glutamic acid. But it is possible that the conservation of the second negative charge and the structural similarities between these two residues enable a robust G protein-antibody interaction. Lastly, VEQ corresponding aa residues in PIRYV.G, VEQ, have electrostatically and structurally different characteristics to that of lysine and aspartic acid leading to the lack of interaction between the mAb and G protein.

We showed that IE9F9 recognises a β-sheet rich domain of the G protein (7, 17). A complete abrogation of binding wasn’t observed with the VSVind.G mutants produced. This implies that the antibody either relies on other structural cues and environmental charges around for binding or can utilise a secondary epitope. However, through the gain of binding effect observed in COCV.G mutants, we were able to identify two regions; AA and LSR, aa residues 352-353 and 356-358 respectively on VSVind.G, that are the key to this antibody’s interaction.

All three reagents investigated demonstrated neutralising activities. 8G5F11 had the greatest ability to cross-neutralise a wide array of vesiculovirus family members. The strength of neutralisation for this mAb, however, didn’t correlate with its affinity towards other VesG (Figure 2 and 3). This suggests that innate differences, such as protein structure, between the VesG might be playing a role in LV neutralisation. Since the structures of the VesG other than VSVind.G and CHAV.G are not yet delineated, it is hard to accurately point out the key factors and mechanism involved. However, we have identified 8G5F11’s epitope to lie close to the cross-over point between pleckstrin homology and trimerisation domain of VSVind.G (7, 17, 19, 20, 35). Several hinge segments have been identified in the proximity of the epitope which undergo large rearrangements in its relative orientation while the G protein refolds from pre to post-fusion conformation in the low-pH conditions of the endosomes following endocytosis (16, 19, 35). It can be hypothesised that 8G5F11 might be hindering this process ultimately preventing viral fusion and infection. As pH-induced conformational changes during viral fusion is a shared characteristic amongst VesG (43), this might be the underlying reason behind 8G5F11’s ability to cross-neutralise VesG-LV.

We have shown that IE9F9 blocks VSVind.G binding to its major receptor LDLR (Figure 6). The crystal structures of VSVind.G in complex with LDLR domains have been recently identified and have shown that VSVind.G can interact with two distinct cysteine-rich domains (CR2 and CR3) of LDLR (35). One of the regions on VSVind.G that is crucial for LDLR CR domain binding lies between amino acids 366-370, only seven amino acids away from the key residues in IE9F9’s epitope. The key residues in this region of VSVind.G are not conserved amongst vesiculoviruses therefore, neither the use of this epitope nor LDLR can be generalised to the other members of the genus, making IE9F9’s epitope and neutralisation mechanism specific to VSVind.G. The lack of cross-reactivity and cross-neutralisation (Figure 1 and 3) displayed by the mAb towards VesG as well as its failure to neutralise COCV.G mutants when its epitope is inserted into the G protein (Figure 5) suggest specific requirement for binding mode between IE9G9 and G proteins to result in neutralisation. Nikolic and colleagues have demonstrated that VSVind.G has specifically evolved to interact with the CR domains of other LDLR family members (35). The other members of the receptor family have already been identified as secondary ports of entry for the virus (33). Complete neutralisation achieved with IE9F9 indicates that the other LDLR family members might be interacting with the same epitope on VSVind.G as well.

Further work on these two identified epitopes regarding their immunodominance in an *in vivo* setting and their detailed characterisation on other VesG from the structure-function point of view may be of interest in the context of host-pathogen interaction and co-evolution. This may also provide the opportunity for modifying VSVind.G to improve G protein-containing advanced therapy medicinal products and VSVind-based vaccine vectors.

## MATERIALS AND METHODS

### Cell culture

In all experiments, HEK293T cells were used. The cell line was maintained in Dulbecco’s Modified Eagle Medium (DMEM) (Sigma-Aldrich, St Louis, MO) supplemented with 10% heat-inactivated foetal calf serum (Gibco, Carlsbad, CA), 2mM L-Glutamine (Gibco), 50 units/ml Penicillin (Gibco), 50μg/ml Streptomycin (Gibco). All cells were kept in cell culture incubators at 37°C and 5% CO_2_.

### Phylogenetic analysis of vesiculovirus and rabies virus G proteins based on amino acid sequences

G proteins of the major vesiculoviruses (VSVind, UniProt Accession Number: P03522, Cocal virus, #O56677, VSVnj, #P04882, Piry virus, #Q85213, Maraba virus, #F8SPF4, VSVala, # B3FRL4, Chandipura virus, #P13180, Carajas virus, #A0A0D3R1Y6, Isfahan virus, # Q5K2K4) as well as the G protein of the Rabies virus (#Q8JXF6), were included in the analysis. The amino acid sequences were aligned using ClustalOmega online multiple sequence alignment tool (EMBL-EPI). The evolutionary analyses were conducted in MEGA7 (44). The evolutionary history was inferred by using the maximum likelihood method based on the Jones-Taylor-Thornton matrix-based model (45). The tree with the highest likelihood is shown with the bootstrap confidence values (out of 100) indicated at the nodes. The tree is drawn to scale, with branch lengths measured in the number of substitutions per site, depicted in the linear scale.

### Plasmids used in experiments

VSVind.G expression plasmids, pMD2.G, and gag-pol expression plasmid p8.91 (46) were purchased from Plasmid Factory (Germany). GFP expressing self-inactivating vector plasmid used in the production of lentiviral vectors was produced in our lab previously (47, 48). pMD2.Cocal.G, COCV.G, expression plasmid was a kindly provided by Hans-Peter Kiem (Fred Hutchinson Cancer Research Center, Seattle, WA). All other VesG envelopes were cloned into this backbone using the restriction enzymes PmlI and EcoRI. Amino acid sequences for VSVnj.G, PIRYV.G, MARAV.G, VSVala.G were retrieved from UniProt. Codon-optimised genes were ordered from Genewiz (South Plainfield, NJ). Unrelated feline endogenous virus RD114 derived RDpro envelope (48) was used as a negative control in several assays.

### Gene transfer to mammalian cells

Single plasmid transfection was used to express VesG on HEK293T cell surface. HEK293T cells were seeded on the day prior to transfection at 4×10^6^ cell per 10cm plate. These cells were transfected by lipofection using FuGENE6 (Promega, Madison, WI) according to the manufacturer’s instructions. The cells were harvested 48h later to be used in various flow cytometry assays.

### Overlapping extension PCR to synthesise VesG chimeras

Phusion High-Fidelity PCR Kit (NEB, Ipswich, MA) was used to perform the PCR reactions. All primers used were obtained from Sigma-Aldrich (Supplementary Table 1). To splice two DNA molecules, special primers were at the joining ends. For each molecule, the first of two PCRs created a linear insert with a 5’ overhang complementary to the 3’ end of the sequence from the other gene. Following annealing, these extensions allowed the strands of the PCR product to act as a pair of oversized primers and the two sequences were fused. Once both DNA molecules were extended, a second PCR was carried out with only the flanking primers to amplify the newly created double-stranded DNA of the chimeric gene.

### Surface plasmon resonance

Analyses were performed using a BIAcore T100 instrument (GE Healthcare). Gth (0.04 mg/mL) and 8G5F11 (0.03 mg/mL) in sodium acetate buffers (10mM, pH 4.5 and 4.0 respectively) were immobilised on a CM5 sensor chip using the amine coupling system according to the manufacturer’s instructions. To measure mAb affinity to VSVind.G, 8G5F11 (MW 155kDa) and IE9F9 (MW 155kDa) were suspended in HBS-EP (0.01M HEPES pH7.4, 0.15M NaCl, 3mM EDTA, 0.005v/v P20) and passed over the immobilised Gth at the indicated concentrations. To measure VesG-LV avidity against 8G5F11, LV preparations were suspended in HBS-EP buffer and passed over the immobilised mAb at indicated titers. The dissociation constants were calculated using BIAevaluation software according to the manufacturer’s instructions. For the competitive binding assay, multiple injections of mAbs at 10μg/mL concentration was performed followed by injection of soluble recombinant LDLR (R&D Systems, Minneapolis, MN) at an identical concentration.

### Use of molecules of equivalent soluble fluorochrome (MESF) system for quantitative flow-cytometry analysis

Quantum Alexa Fluor 647 MESF kit (Bangs Laboratories, Fishers, IN) was utilised for all quantitative fluorescence flow cytometry experiments. This is a microsphere kit that enables the standardisation of fluorescence intensity units. Beads with a pre-determined number of fluorophores are run on the same day and at the same fluorescence settings as stained cell samples to establish a calibration curve that relates the instrument channel values (i.e. median fluorescence intensity (MFI)) to standardised fluorescence intensity (MESF) units.

### SDS/PAGE

Gth was visualised via Ponceau S staining. 15μg of Gth was boiled at 95°C for 5 mins in 5X Laemmli buffer (5% Sodium dodecyl sulfate (SDS), 50% glycerol, 0.1% bromophenol blue, 250mM Tris-HCl, pH 6.8, and, 5% β-mercaptoethanol) and resolved on 10% SDS-PAGE gel (10% acrylamide-Tris). Sample was then transferred onto a nitrocellulose membrane (GE Healthcare) and visualised.

### Extracellular and intracellular antibody binding assay

HEK293T cells were transfected to express the G proteins. 48 hours later cells were harvested, washed twice with PBS and plated in U-bottom 96-well plates at identical densities. For intracellular antibody binding assays cells were fixed with 1% formaldehyde (Sigma-Aldrich, St Louis, MO) in PBS, permeabilised using 0.05% saponin (Sigma-Aldrich, St Louis MO) in PBS and blocked with 1% bovine serum albumin (BSA, Sigma-Aldrich, St Louis MO) in PBS. Cells were then incubated with serial dilutions of extracellular and intracellular antibodies ranging from 0.1mg/ml to 2×10-7 mg/ml in 1% BSA (Sigma) in PBS in a total reaction volume of 200μl. After washing twice, each sample was incubated with its respective fluorophore-conjugated secondary antibody (Antibodies used are listed in Supplementary Table 2). Cells were then washed twice and resuspended in PBS. Stained cell samples were analysed via flow cytometry using a FACSCanto II (BD Biosciences, San Jose, CA) and Flowjo software.

### Transient LV production and concentration

Three-plasmid co-transfection into HEK293T cells was used to make pseudotyped LV as described previously (46). Briefly, 4×10^6^ 293T cells were seeded in 10cm plates. 24 hours later, they were transfected using FuGene6 (Promega, Madison, WI) with following plasmids: SIN pHV (GFP expressing vector plasmid (47, 48)), p8.91 (Gag-Pol expression plasmid (46)), and envelope expression plasmids. The medium was changed after 24 hours and then vector containing media (VCM) was collected over 24-hour periods for 2 days. Following collection, VCM was passed through Whatman Puradisc 0.45μm filters (SLS) and concentrated ∼100-fold by ultra-centrifugation at 22,000 rpm (87,119xg) for 2 hours at 4°C in Beckmann Optima LK-90 ultracentrifuge using the SW-28 swinging bucket rotor (radius 16.1cm). The virus was resuspended in cold plain Opti-MEM on ice, aliquoted and stored at −80°C.

### LV titration

The functional titre of each vector preparation was determined by flow cytometric analysis for GFP expression following transduction of HEK293T cells. Briefly, 2×10^5^/well 293T cells were infected with LV plus 8 μg/ml polybrene (Merck-Millipore, Billerica, MA) for 24 hours. Infected cells were detected by GFP expression at 48 hours following the start of transduction. Titres were calculated from virus dilutions where 1–20% of the cell population was GFP-positive using the following formula:

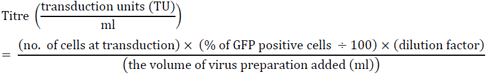

### Antibody neutralisation assay

To determine the neutralisation activity of anti-VSVind.G monoclonal and polyclonal antibodies an infection assay in the presence of antibodies was performed. Briefly, HEK293T cells were seeded in a 96-well plate at a density of 2×10^4^ cells/well with 200μl of medium containing 8μg/ml polybrene. Approximately 3 hours later, antibodies were serially diluted in plain Opti-MEM to 12 different concentrations/dilutions ranging from 0.5mg/ml (1:2 dilution) to 1.6×10^−7^ mg/ml (1:6,250,000 dilution). Each antibody dilution was mixed 1:1 with VesG-LV or mutant G-LV at 4.0×10^5^TU/ml titre to a final volume of 20μl, incubated at 37°C for 1h and plated on the cells. 48 hours after cells were harvested and analysed for GFP expression by flow cytometry.

### Site-directed mutagenesis PCR for production of mutant G proteins for epitope mapping

Site-directed mutagenesis (SMD) method was utilized to produce G protein mutants that were used in epitope mapping experiments. For this, QuikChange II XL Site-Directed Mutagenesis Kit (Agilent, Santa Clara, CA) was used. Initially, primers that would have the desired nucleotide changes were designed using the QuikChange Primer Design Tool (http://www.genomics.agilent.com/primerDesignProgram.jsp). All primers used were obtained from Sigma-Aldrich (St Louis, MO). The reaction was carried out according to manufacturer’s instructions.

## Funding

AMM and MT studentships are funded by NIBSC.

### Acknowledgements

We would like to thank Dr Hiroo Hoshino and Dr Atsushi Oue (Gunma University, Japan), for providing us with a sample of the polyclonal goat anti-VSVind.G antibody, VSV-Poly and Drs Yves Gaudin and Aurélie Albertini for the Gth protein.

## Author Contributions

AMM performed experiments to obtain presented data and wrote the paper. MT designed and produced the initial COCV.G/VSVind.G chimeras and obtained preliminary data on 8G5F11 binding to COCV.G bearing cells. GM and MH helped designing experiments and interpreting data. MKC and YT supervised the study and wrote the paper.

## Additional Information

### Supplementary information

accompanies this manuscript.

### Competing financial interest

Authors declare no competing financial interests.

## Footnotes

1 It should be noted that the amino acid sequence of the full-length G proteins (including the signal peptide) were referred to in this manuscript. Accordingly, reference to specific residue numbers is made in the context of these full-length sequences.

